# The enemy of my enemy is my friend: a plausible defense mechanism against antagonism in bacterial communities

**DOI:** 10.1101/2023.04.17.537116

**Authors:** Laura Sánchez-Gómez, Moisés Santillán

## Abstract

In a recent work we studied an artificial bacterial community where resistant bacteria assisted sensitive bacteria in surviving in the presence of antagonistic bacteria. Typically, this behavior is attributed to lateral gene exchange. However, we identified evidence of a distinct mechanism, which operates on the principle of “the enemy of my enemy is my friend”. Here, we explore the viability of this mechanism by means of a reaction-diffusion mathematical model. Our findings suggest that this mechanism is feasible under specific circumstances, particularly when bacteria undergo significant metabolic changes at high population densities.

**Author summary:** The importance of studying the formation of bacterial communities cannot be overstated. Initially, it was believed that the environment alone determined the assembly of these communities, but recent research has demonstrated that interactions among members of the community also have a crucial impact. For instance, it has been observed in some cases that, even when susceptible bacteria are confronted with antagonism, they can survive with the aid of resistant bacteria. Lateral gene exchange is typically considered the underlying mechanism responsible for this phenomenon. However, we identified in a recent work an alternative mechanism that does not involve gene exchange but operates on the principle of “the enemy of my enemy is my friend”. Using a mathematical model, we examine the plausibility of this mechanism in a small artificial bacterial community. Our results indicate that this mechanism alone cannot account for all the experimental observations, but including the assumption that bacteria undergo significant metabolic changes at high concentrations may suffice.

## Introduction

Bacteria, which have been present on Earth for billions of years, are capable of thriving in even the most hostile environments [1, 2]. However, their intriguing traits are not limited to their ability to survive almost anywhere. The interactions between bacteria are also critical for the existence of complex communities in which multiple species can compete for or share resources, allowing them to thrive [3–5]. These communities are essential for biosphere sustainability, as they are responsible for important processes such as decomposition of dead organisms and nitrogen fixation [2, 6, 7].

Past studies on microorganisms have primarily focused on the physiology and growth of bacteria in pure culture rather than their interactions and formation of communities [1, 8–11]. However, there has been a recent surge in interest in the significance of bacterial communities [2, 4, 5, 12–17]. This area of research addresses two major concerns: 1) the factors that contribute to bacterial biodiversity [2, 12, 18–22], and 2) the mechanisms that promote the formation of spatial patterns within these communities [9, 12, 17, 23–30].

It was previously believed that the formation of communities was solely determined by environmental factors [31]. Consequently, abiotic interactions have received more attention than biotic interactions [32–37]. However, bacteria are highly social organisms that can form complex communities with high levels of biodiversity [12]. These communities are the result of both abiotic and biotic interactions, including responses to environmental cues (such as nutrient availability, pH, and physical space) and interactions with neighboring cells of the same or different strains [12, 38–41].

When it comes to biotic interactions, bacteria have developed various adaptable strategies to ensure population survival within communities. These strategies include altering their metabolism, working together, competing, and even programmed death. Competition encompasses a wide range of behaviors, including competing with individuals of the same and different strains [3] and antagonism [12, 19, 42, 43].

Specifically, the role of antagonism, which involves the release of harmful substances like antibiotics or toxins that diffuse in the medium and hinder or eliminate competitors, has been established as central in bacterial life [43].

Bacterial strains can reap benefits from coexisting with other strains. For instance, drug resistance can be transferred between strains, and some resistant bacteria can assist sensitive bacteria in surviving under the presence of antagonistic strains [44–46]. For the sake of brevity, here and thereafter we refer to this latter phenomenon as “transfer of resistance against antagonism”.

The Churince lagoon is a part of an isolated aquifer body located in the middle of the Cuatro Ciénegas valley in Mexico. The valley is made up of several pools and streams that are connected underground through a network of faults resulting from the geological activity in the region [47]. From the sediment of the Churince lagoon, Cerritos *et al*. [21] recently isolated a group of thermoresistant bacteria. The collection consisted of 78 strains gathered from various sampling sites throughout the lagoon, and later Pérez-Gutiérrez *et al*. [29] recorded over 6000 antagonistic interactions among them. Due to the high occurrence of antagonistic interactions and the physicochemical characteristics of the lagoon, the Churince isolates are suitable candidates for investigating the transfer of resistance against antagonism.

Two separate investigations were carried out to examine the transfer of resistance against antagonism in simplified bacterial communities using bacterial strains derived from the Churince lagoon [21]. In the first study conducted by Gallardo-Navarro and

Santillán [44], *Bacillus pumilus* (antagonistic), *Bacillus aquamaris* (sensitive), and *Staphylococcus sp*. (resistant) were utilized, while the second study by Aguilar-Salinas and Olmedo-Álvarez [45] employed *Bacillus pumilus* (antagonistic), *Sutclifiella horikoshii* (sensitive), and *Bacillus cereus* (resistant). Both studies established that transfer of resistance against antagonism can occur in vitro in simplified bacterial communities. However, while Aguilar-Salinas and Olmedo-Álvarez [45] proposed that the responsible mechanism is horizontal gene exchange, Gallardo-Navarro and SantillÁn [44] argued that gene exchange cannot account for their findings and suggested an alternative mechanism that operates on the principle of “the enemy of my enemy is my friend”.

This study aims to explore the mechanism proposed by Gallardo-Navarro and Santillán [44] for transfer of resistance against antagonism. According to this proposal, antagonistic bacteria can sense the presence of other bacteria through a hypothetical metabolite produced by the three of them and respond by producing an antagonistic substance, at a metabolic cost. The antagonistic substance can kill the sensitive strain, while the resistant strain can counteract it, but this comes at the expense of decreasing their growth rate. In the presence of resistant bacteria, antagonistic bacteria produce more antagonistic substance, which slows the growth of sensitive bacteria, thereby indirectly facilitating their survival. To assess the feasibility of this machinery, we employ mathematical modeling based on reaction-diffusion equations.

## Methods

### Mathematical model development

Mathematical models can be broadly classified into two categories: predictive and explanatory models. Predictive models are designed to generate precise quantitative predictions, while explanatory models focus on identifying causal relationships between input variables and outcomes, even if their predictions may not be entirely accurate. In this section, we will focus on developing an explanatory mathematical model to investigate the bacterial interaction mechanisms proposed by Gallardo-Navarro and SantillÁn [44]. The model will consider a community consisting of three bacterial strains: an antagonist (*A*), a resistant (*R*), and a sensitive (*S*) strain. The model assumes that the antagonist strain can sense a metabolite (*m*) produced by all three bacterial strains and, in response, produces an antagonistic substance (*u*). The production of the antagonistic substance incurs a metabolic cost for the antagonist strain, while the resistant strain pays a metabolic cost to resist the substance, and the sensitive strain is killed by it. The above-described interactions are represented mathematically by a system of reaction-diffusion partial differential equations (PDEs):

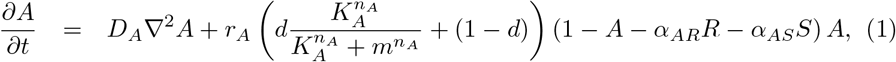

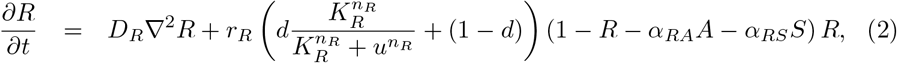

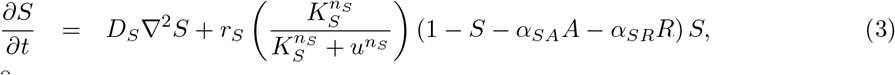

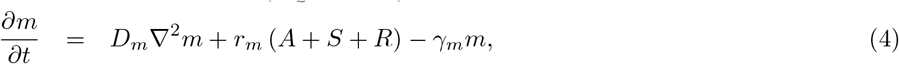

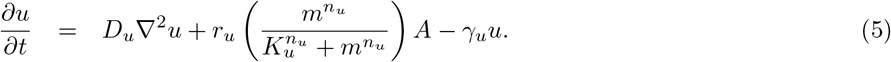

The equations above pertain to the concentration of bacterial strains or chemical species at a given location and time, where the bacterial strain variables are normalized to their respective carrying capacity. Each equation’s right-hand side has a diffusion term, and *D*_*i*_ denotes the diffusion coefficient for each species (*i* can be *A, R, S, m*, or *u*). The subsequent term(s) indicate the growth/production and death/decay rates for each species, with *r*_*i*_ representing the growth/production rate for *i* = *A, R, S, m*, or *u*, while *γ*_*m*_ and *γ*_*u*_ denote the decay rates for the metabolite and antagonistic substances, respectively.

The present model was constructed to mimic the experimental conditions of Gallardo-Navarro and Santillán [44], in which the growing medium was abundant in nutrients. Therefore, bacterial growth in single-strain cultures was assumed to be logistic. Additionally, Gallardo-Navarro and Santillán [44] observed that resistant bacteria coexisted with antagonistic and sensitive strains, while the latter two were mutually exclusive. To capture these dynamics, we utilized Lotka-Volterra competition equations to model the growth of bacterial communities. Here, *α*_*ij*_ represents the effect of species *j* on the growth rate of species *i*, where *i, j* = *A, R* or *S*.

The bacterial strains used in the study by Gallardo-Navarro and SantillÁn [44] have not been fully characterized, and therefore, it is not possible to provide detailed information regarding the mechanisms underlying the interactions with the hypothesized antagonistic substance and common metabolite. However, it is likely that cooperative ligand-receptor interactions are involved. Based on this, we assume that Hill-like functions can be used to model the response functions that account for the metabolic cost of producing or counteracting the antagonistic substance, as well as its toxic effect on sensitive bacteria [48].

Eq (1) models the dynamics of the antagonistic strain (*A*), whose growth rate (*r*_*A*_) is modulated by a decreasing Hill-like function of the concentration of metabolite (*m*).

This modulation stands for the metabolic cost associated to the synthesis and release of the antagonistic substance, in response to the presence of *m*. Notice that the associated growth rate decrease is bounded by parameter 0 ≤ *d* ≤ 1 (when *d* = 0, high concentrations of *m* can completely halt the growth of *A*, whereas when *d* = 1, the growth of *A* remains unaffected by *m*). This accounts for the fact that, as observed by Gallardo-Navarro and SantillÁn [44], the growth of the antagonistic strain is not totally stopped when it synthesizes the antagonistic substance. *K*_*A*_ is the concentration of *m* needed to reduce *r*_*A*_ down to *r*_*A*_ (1 − *d/*2), and *n*_*A*_ is a Hill coefficient.

Eq (2) corresponds to the dynamics of the resistant strain (*R*). Its growth rate (*r*_*R*_) is regulated by a decreasing Hill-like function of the antagonistic substance (*u*). This represents the ability of *R* to resist *u*, which comes with a metabolic cost as well. *K*_*R*_ accounts for the concentration of *u* needed to reduce *r*_*R*_ down to *r*_*R*_ (1 − *d/*2), and *n*_*R*_ is a Hill coefficient. Analogous to the case of *A*, the presence of *u* is not enough to stop *R*’s growth completely [44], hence the presence of parameter *d*.

Eq (3) represents the dynamics of the sensitive strain (*S*). Its growth rate (*r*_*S*_) is regulated by a decreasing Hill function of the antagonistic substance (*u*). Given that *u* kills and prevents *S* from growing, the strain growth rate can become zero [44]. *K*_*S*_ is the concentration of *u* needed to reduce *r*_*S*_ to half its maximal value, and *n*_*S*_ is a Hill coefficient.

Eq (4) stands for the dynamics of the metabolite (*m*). Metabolite production (*r*_*m*_) is proportional to the total bacterial concentration, considering all strains; whereas decay is assumed exponential, with rate constant *γ*_*m*_.

Finally, Eq (5) denotes the dynamics of the antagonistic substance (*u*). An increase in the local concentration of *m* initiates the production of *u* by antagonistic bacteria (notice that antagonistic substance production rate is proportional to *A*), which is represented by an increasing Hill function of *m* with maximum rate *r*_*u*_. *K*_*u*_ is the half saturation constant and *n*_*u*_ is a Hill coefficient. As in *m*, the decay of *u* is assumed exponential, with rate constant *γ*_*u*_.

It is important to mention that all the parameters of the model have positive values. Because of this and the fact that *d* is less than or equal to 1, standard comparison principles imply that if the initial data is bounded and non-negative, then *A, R*, and *S* will also be bounded. In fact, if the initial values satisfy 0 ≤ *A*(*x*, 0), *R*(*x*, 0), *S*(*x*, 0) ≤ 1 then 0 ≤ *A*(*x, t*), *R*(*x, t*), *S*(*x, t*) ≤ 1 for all *t >* 0, and consequently, if follows from the equation for *m* that *m* is bounded, which implies from the equation for *u* that it is also bounded. Furthermore, even if the initial values for *A, R, S* are not bounded above by 1, it can still be inferred that non-negative initial data in all components leads to:

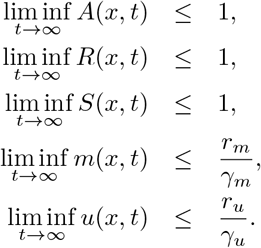

### Numerical solution of the model PDE system

The model PDE system was numerically solved on bounded rectangular or square areas corresponding to the surface on which bacterial populations grow. As the model is employed to replicate Petri dish experiments and the simulated region represents a portion of the agar surface far enough from the dish boundaries, Dirichlet boundary conditions were employed.

Regular square lattices were constructed within the simulated surfaces. The discretization of the Laplacian was performed in every compartment of the lattice, using a fourth order 9-point stencil as described in [49]:

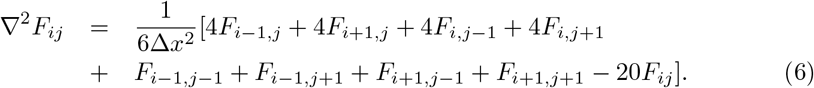

In this equation, *F*_*i,j*_ represents the average value of function *F* in the compartment with coordinates *i, j*, ▽^2^*F*_*i,j*_ denotes the Laplacian of *F* in the same compartment, and Δ*x* represents the length of the lattice-compartment side. Its value was set to Δ*x* = 0.1 mm and the time period corresponding to one algorithm iteration was set to Δ*t* = 0.01 min. These values were chosen by empirically testing the algorithm numerical convergence.

Since we considered Dirichlet boundary conditions, the computation of the Laplacian involved three types of lattice compartments based on their location and the number of adjacent compartments: interior, edges, and vertices. We employed Eq (6) to calculate the Laplacian for interior compartments, while for edges or vertices, we adjusted the equation to account for the function *F* taking zero values in compartments beyond the simulation lattice.

Taking the above considerations into account, we implemented Euler’s explicit algorithm in Python, to solve the model PDE system in the discretized surface. In numerical analysis, it is generally agreed upon that explicit methods are not very efficient for solving diffusion problems. Therefore, to avoid inaccuracies, we conducted empirical tests to ensure the numerical convergence of the algorithm, carefully selecting values for Δ*x* and Δ*t*. The algorithm pseudo-code is as follows:

1. Set the initial values of variables *A, S* and *R, m* and *u* in all lattice compartments.
2. Calculate the Laplacian of each variable in every compartment of the lattice. The resulting value, when multiplied by the corresponding diffusion coefficient and the time step Δ*t*, determines the change of the variable in each compartment due to diffusion in one iteration.
3. Compute the production and/or degradation of all model variables in each lattice compartment based on the differential-equation terms. These values multiplied by Δ*t* represent the quantity of each variable that is added or removed in each compartment due to the reaction terms of the equations in one iteration.
4. Update the values of all variables by taking into account the changes caused by the reaction and diffusion terms of their respective equations.
5. Iterate to step two.

The Python code is available for interested readers in the repository located at https://github.com/lsanchezg89/facilitationbacterialcommunity.git.

### Intrinsic growth rate estimation

Gallardo-Navarro and SantillÁn [44] conducted bacterial growth experiments in a liquid medium where they measured bacterial concentration periodically for several hours. In order to estimate the intrinsic bacterial growth rates, the reported data was normalized to the highest concentration of each bacterial strain, and then fitted to the following logistic-model function, using the `curve_fit` algorithm from the `SciPy.Optimize` library in `Python`.

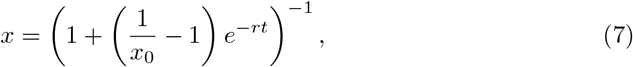

where *x*_0_ denotes the normalized initial bacterial concentration. The best-fitting *r* value can be interpreted as the intrinsic growth rate of the corresponding bacterial strain.

### Single colony simulations

Gallardo-Navarro and SantillÁn [44] performed experiments in which single colonies were inoculated on Petri dishes with marine medium plus 2% agar, and their radii were monitored over a period of 6 days. To simulate these experiments we considered a

45-compartment long square lattice. Since the simulated areas correspond to a section of the agar that is far away from the dish walls, we used Dirichlet boundary conditions. A circle-like area with 4-compartment long radius was identified at the lattice center (see Fig. 1A). In each of the compartments of such circle, the initial value of the variable denoting the concentration of the simulated bacterial strain was randomly selected from a normal distribution with mean 0.2 and standard deviation 0.02.

**Fig 1.**
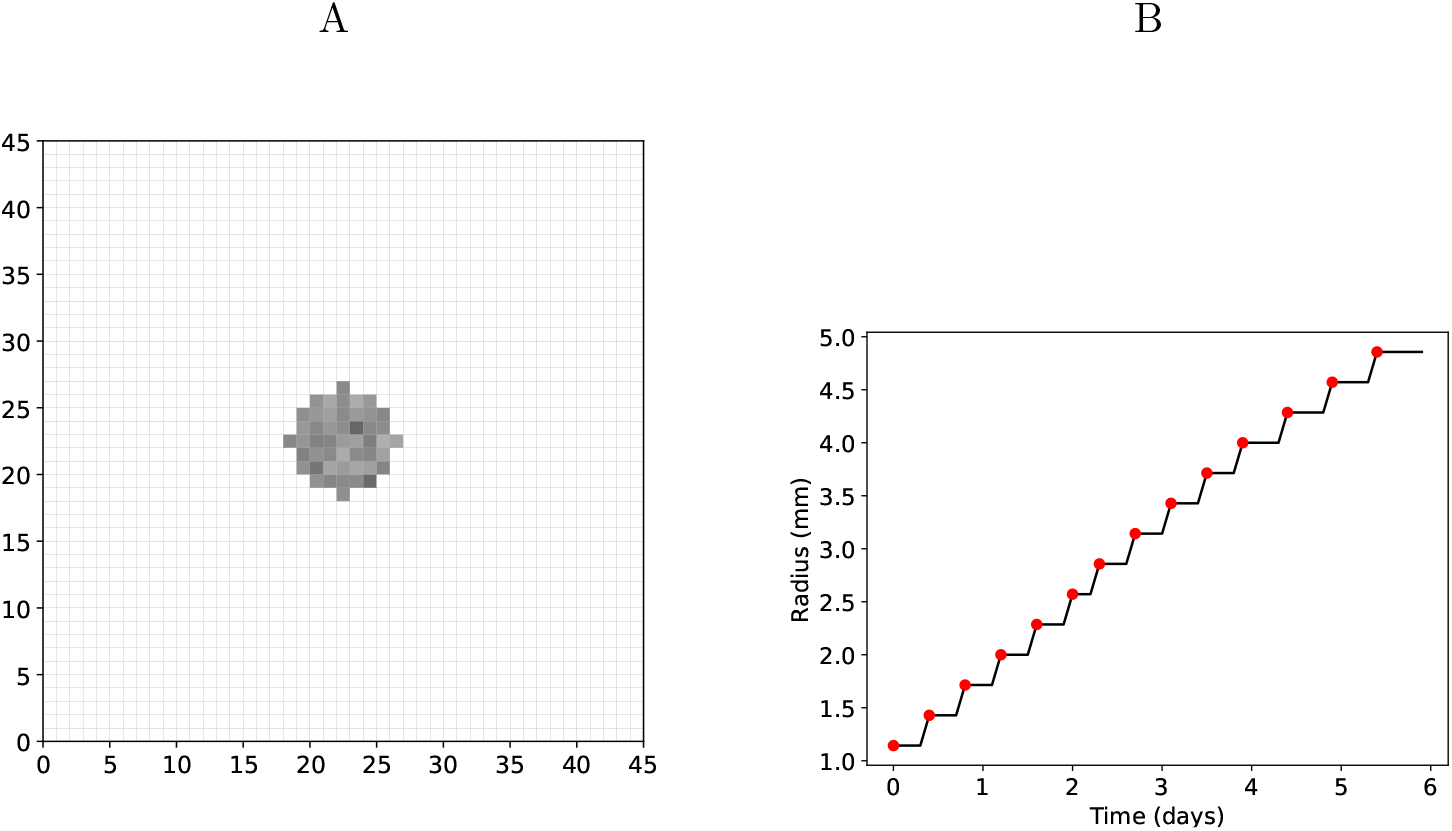
A) Illustration of the initial-condition selection in single-colony simulations. B) Illustration of the radius measurement results in single-colony simulations (solid line). The stair vertices used for interpolation are emphasized (solid red points).

Elsewhere, the bacterial initial concentration was set to zero. The initial values of the variables corresponding to the concentrations of other bacterial populations and metabolites *m* and *u* were set to zero everywhere. The spatiotemporal evolution of the colony was simulated by solving the model PDE set as previously described. The simulation results were carried out until the simulation time reached 6 days and the results were recorded every simulated minute.

To measure the colony radius at a given time, we first identified compartments with bacterial concentrations less than 0.01 and set their values to zero. Then, we counted the number of bacteria containing compartments along the colony diameter, and computed the radius by dividing the result by 2 and multiplying by Δ*x*. Due to the spatial discretization inherent to the numerical method used to solve the model equations, the obtained radius vs. time functions resulted to be step-like (see Fig. 1B). To get smooth curves from them, we took the step vertices and performed an interpolation with algorithm interp1d from Python’s library SciPy.Interpolate.

### Simulations of confronted colonies

Gallardo-Navarro and SantillÁn [44] carried out experiments with confronted colonies in which a couple of 1 μl bacterial-culture drops, of the same or different strains, were inoculated with a separation of 15 mm on a Petri dish with marine medium plus 2% agar; and the external (in the direction opposite to the other colony) and internal (in the direction of the other colony) radii were measured periodically over a period of 6 days. To simulate these experiments, we used 90 × 144 rectangular lattices. Lattices represent an area of the agar sufficiently far away from the dish walls, and hence Dirichlet boundary conditions were used. Drop inoculation was mimicked by setting the initial values of the variables corresponding to the inoculated strains equal zero everywhere, except in circle-like areas with 4-compartment long radii. These areas are separated by 53 compartments (see Fig. 2). Initial bacterial concentrations were randomly selected from a normal distribution with mean 0.2 and standard deviation 0.02, in each of the compartments within the inoculated areas. The initial values of all other variables were set to zero everywhere. Model equations were numerically solved using the previously described numerical method. They were extended up to a simulation time of 10 days and the simulation results were recorded every simulated minute. Using a procedure similar to the one described in the previous subsection, we measured the external and internal radii of both colonies (see Fig. 2) at all recorded times.

**Fig 2.**
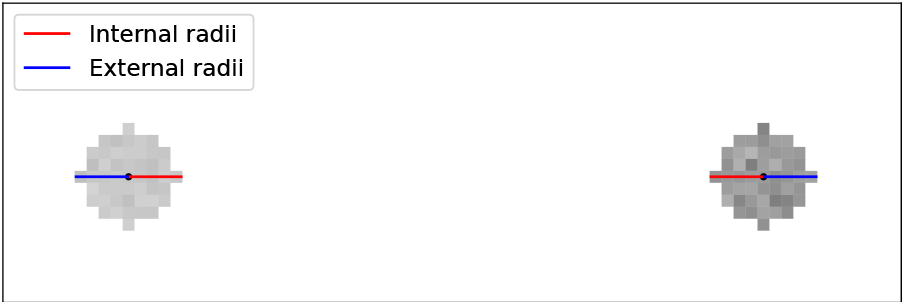
Illustration of the initial-condition selection, and of the internal and external radii measurement, for the simulations of two confronted colonies.

### Artificial-community simulations

Gallardo-Navarro and Santillán [44] conducted two types of experiments involving artificial communities. In the first one they created artificial bacterial communities by mixing a fixed amount of sensitive bacteria with different amounts of antagonists: 10, 25, 50, 75, 100 and 200% of the initial concentration of sensitive bacteria. 5 μl of the mixture was spread on a Petri dish with marine medium plus 2% agar, and incubated for 4 days. The second experiment consisted in adding to the mixtures previously described a fixed amount (500%) of resistant bacteria. In both experiments the authors measured the final area-fraction occupied by sensitive bacteria in terms of the initial antagonistic bacterial concentration.

To simulate these experiments we considered a 60 × 60 square lattice. We used Dirichlet boundary conditions, given that the simulated areas correspond to a region of the agar that is sufficiently far from the dish walls. The initial concentration of sensitive bacteria was randomly selected in each lattice compartment from a normal distribution with mean 0.2 and standard deviation 0.02. The initial concentration of antagonistic bacteria was randomly selected in each lattice compartment from a normal distribution with mean value varying as in the experiments, and standard deviation of 10% of the corresponding mean value. For the first type of experiments, the initial concentration of resistant bacteria was set to zero everywhere, whereas for the second type of experiments, it was randomly selected in each lattice compartment from a normal distribution with mean 1.0 and standard deviation 0.1. The initial values of all other variables were set to zero everywhere.

The model PDE system was then numerically solved up to a simulation time of 20 days and we recorded the simulation results every simulated day. To contrast with experimental results, we computed the fraction of compartments occupied by the sensitive strain at each recorded time.

## Results

Gallardo-Navarro and SantillÁn [44] performed experiments with an artificial community of wild-type bacterial (an antagonistic, a sensitive, and a resistant strain) and found that the presence of resistant bacteria aids the sensitive ones in surviving. They also demonstrated that the previously proposed mechanism of resistant bacteria spatially isolating the sensitive from the antagonistic ones [30] does not work for the strains they worked with, and instead proposed the following:

- By sensing an increase in the concentration of a common metabolite that diffuses in the medium, antagonistic bacteria detect the presence of other bacteria.
- In response, antagonists begin the production and secretion of an antagonistic substance in order to reduce competition, at the expense of metabolic cost.
- This antagonistic substance kills as well as inhibits the growth of sensitive bacteria.
- Resistant bacteria can counteract the antagonistic-substance effect, albeit at a metabolic cost.
- The growth of antagonistic bacteria is slowed down by resistant bacteria, by inducing them to produce the antagonistic substance. This, in turn, indirectly promotes the growth of sensitive bacteria.

The proposed mechanism by Gallardo-Navarro and SantillÁn [44] were supported by indirect evidence, but it was deemed insufficient. Therefore, we constructed a reaction-diffusion PDE model in this study that includes all of the suggested interactions to determine if they suffice to explain the observed behavior or if adjustments are necessary.

To achieve our goal, we tackled the inverse problem by seeking parameter values that would enable the model to replicate the experiments conducted by Gallardo-Navarro and SantillÁn [44]. Instead of attempting to fit all experimental data sets simultaneously, we began with experiments that could be simulated using simpler model versions. This enabled us to estimate a subset of parameter values. We then moved on to more complex experiments and repeated the process until we arrived at experiments that required the complete model. Working in this manner had the advantage of dealing with fewer parameters at a time. However, we discovered that the problem was a challenging multi-objective optimization problem because no single parameter setting could enable the model to fit all experiments. Then, we fine-tuned all parameter values at the end to obtain a reasonable fit to the experimental data sets. This explains why the accuracy of individual fits may not be excellent. With the exception of the estimation of bacterial intrinsic growth rates, which was achieved through a standard optimization algorithm, we relied on empirical methods to fit the model to all other experimental data sets. The reasons for this approach will become apparent as we discuss the results obtained.

The initial step involved fitting the logistic model’s growth function to the normalized-population versus time data sets of bacterial growth experiments in liquid medium from Gallardo-Navarro and SantillÁn [44]. The outcomes are presented in Figs. 3A-C. From these fits, we were able to determine the intrinsic growth rate constants for the antagonistic, resistant, and sensitive strains (*r*_*A*_, *r*_*R*_ and *r*_*S*_, respectively) that are listed in Table 1. Observe that the resistant strain grows much more slowly than the antagonistic and sensitive strains. This result is consistent with prior observations [44].

**Fig 3.**
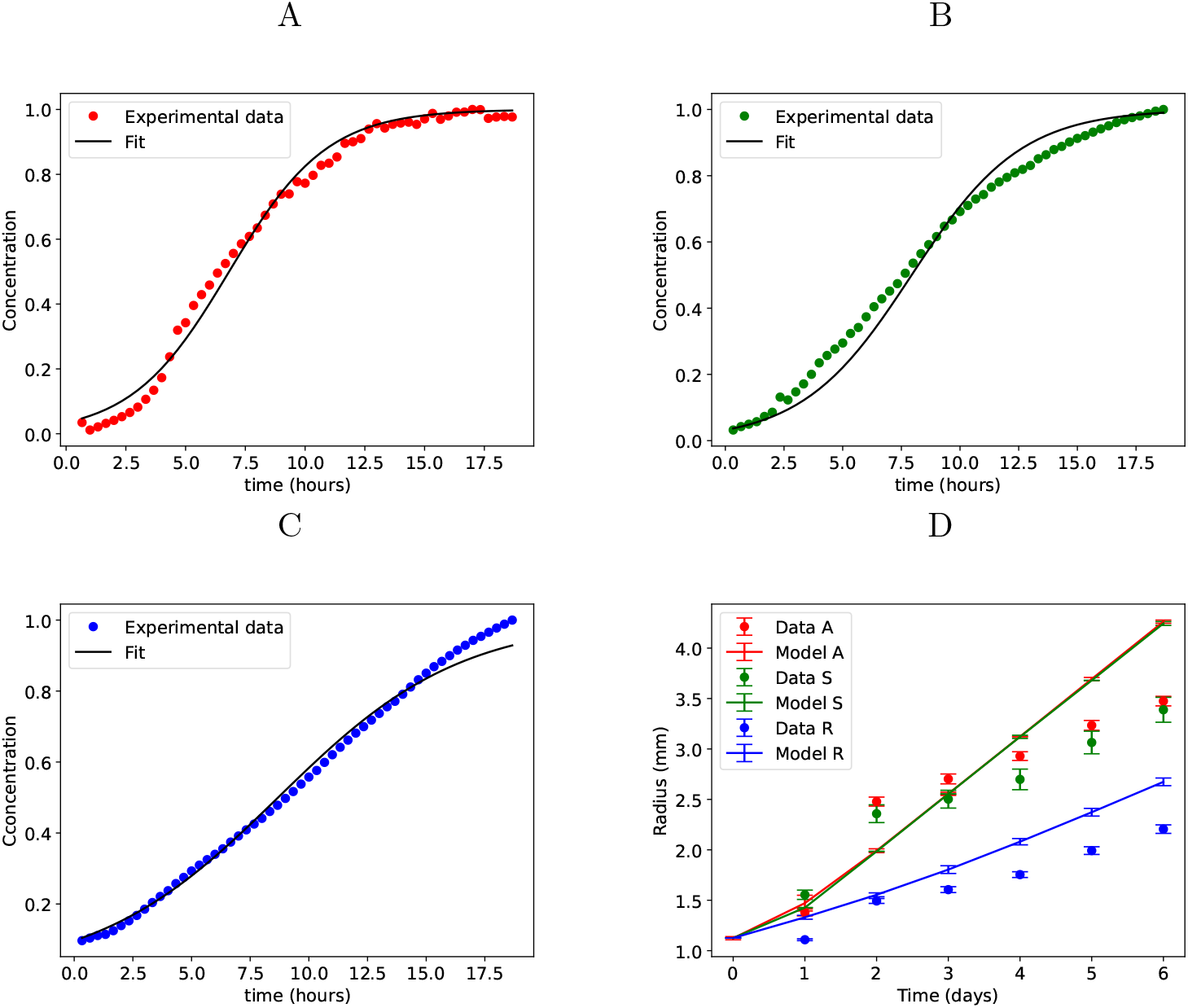
Experimental growth curves [44] and best fit to logistic-model equations for the antagonistic (A), sensitive (B) and resistant (C) strains. D) Comparison of the simulations of single colonies growing on agar (solid lines) and the corresponding reported experimental results (points). Plots represent the average of five simulations and error bars the corresponding standard deviation.

**Table 1.** Model parameter values for single-colony and confronted-colonies simulations.

| | $A$ | $R$ | $S$ | $m$ | $u$ |
| --- | --- | --- | --- | --- | --- |
| $D$ (mm<sup>2</sup>/min) | $1.7 \times 10^{-7}$ | $5.1 \times 10^{-8}$ | $1.02 \times 10^{-7}$ | 0.34 | 0.1615 |
| $r$ (min<sup>-1</sup>) | 0.0078 | 0.0047 | 0.0086 | 0.2 | 0.2 |
| $K$ | $1.0 \times 10^{-8}$ | $1.0 \times 10^{-28}$ | $1.0 \times 10^{-14}$ | NA | $1.0 \times 10^{-12}$ |
| $n$ | 3.5 | 3.0 | 3.5 | NA | 3.5 |
| $\gamma$ (min<sup>-1</sup>) | NA | NA | NA | 0.01 | 0.01 |
| $d$ | 0.25 | 0.25 | NA | NA | NA |

**Table 2.** Estimated values for close-range competition parameters, which are employed in the artificial-community simulations.

| Parameter | Value |
| --- | --- |
| $\alpha_{AS}$ | 2.0 |
| $\alpha_{SA}$ | 1.65 |
| $\alpha_{AR}$ | 0.5 |
| $\alpha_{RA}$ | 0.6 |
| $\alpha_{SR}$ | 0.6 |
| $\alpha_{RS}$ | 0.4 |

Subsequently, we proceeded to estimate the diffusion coefficients for the three bacterial strains. For this purpose, we utilized our mathematical model to simulate the single colony experiments conducted by Gallardo-Navarro and Santillán [44]. We used the previously determined intrinsic growth rate constants and adjusted the corresponding diffusion coefficients through empirical means to fit the reported data on colony radius vs. time. The resulting values of *D*_*A*_, *D*_*R*_, and *D*_*S*_ are presented in Table 1, and their corresponding fitting outcomes are displayed in Fig. 3D. Observe that the diffusion coefficients of the antagonistic and sensitive strains are much higher than those of the resistant strain. We explain this result by the fact that the resistant strain is not motile, whereas the sensitive and antagonistic strains are. Moreover, it should be noted that the model can only approximately replicate the experimental growth curves.

Although the experimental growth curves exhibit sigmoidal behavior (particularly those for *A* and *S*), the simulated curves continue to grow linearly at long times. We posit that this is due to bacterial movement on agar plates, which involves swarming rather than Brownian motion, making it difficult to accurately model using traditional diffusion [50]. Consequently, we selected diffusion coefficient values that provided satisfactory fits for short times (less than 3 hours).

We then shifted our focus towards the experiments conducted by Gallardo-Navarro and Santillán [44], which involved confronting an antagonist colony with either an antagonist, sensitive, or resistant colony. Our objective was to estimate the parameters in the Hill functions that account for the interactions with the common metabolite and the antagonistic substance. As described in section Methods, we numerically solved the PDE system given by Eqs. (1)-(5) to simulate these experiments. The diffusion coefficient for the hypothetical antagonistic substance (*D*_*u*_) was estimated (see Table 1) by considering the reported diffusion coefficients of chemoattractants in agar [51–53]. Due to the fact that all three of the bacterial strains in [44] were aerobic, we chose a slightly higher value for *D*_*m*_, considering carbon dioxide as a potential common metabolite (see Table 1). Finally, we took *K*_*A*_, *K*_*R*_, *K*_*S*_, *K*_*u*_, *n*_*A*_, *n*_*R*_, *n*_*S*_, *n*_*u*_ and *d* (*i*.*e*. the parameters in the Hill and Hill-like functions in Eqs. (1) to (5)) as free parameters and empirically changed their values to fit the experimental results reported in [44]. For each simulated situation we recorded the time evolution of the external and internal radii of both confronted colonies (see section Methods) and computed the difference. The outcomes of the best fits we could get are shown in Figs. 4A-C. The resulting parameter values are listed in Table 1.

**Fig 4.**
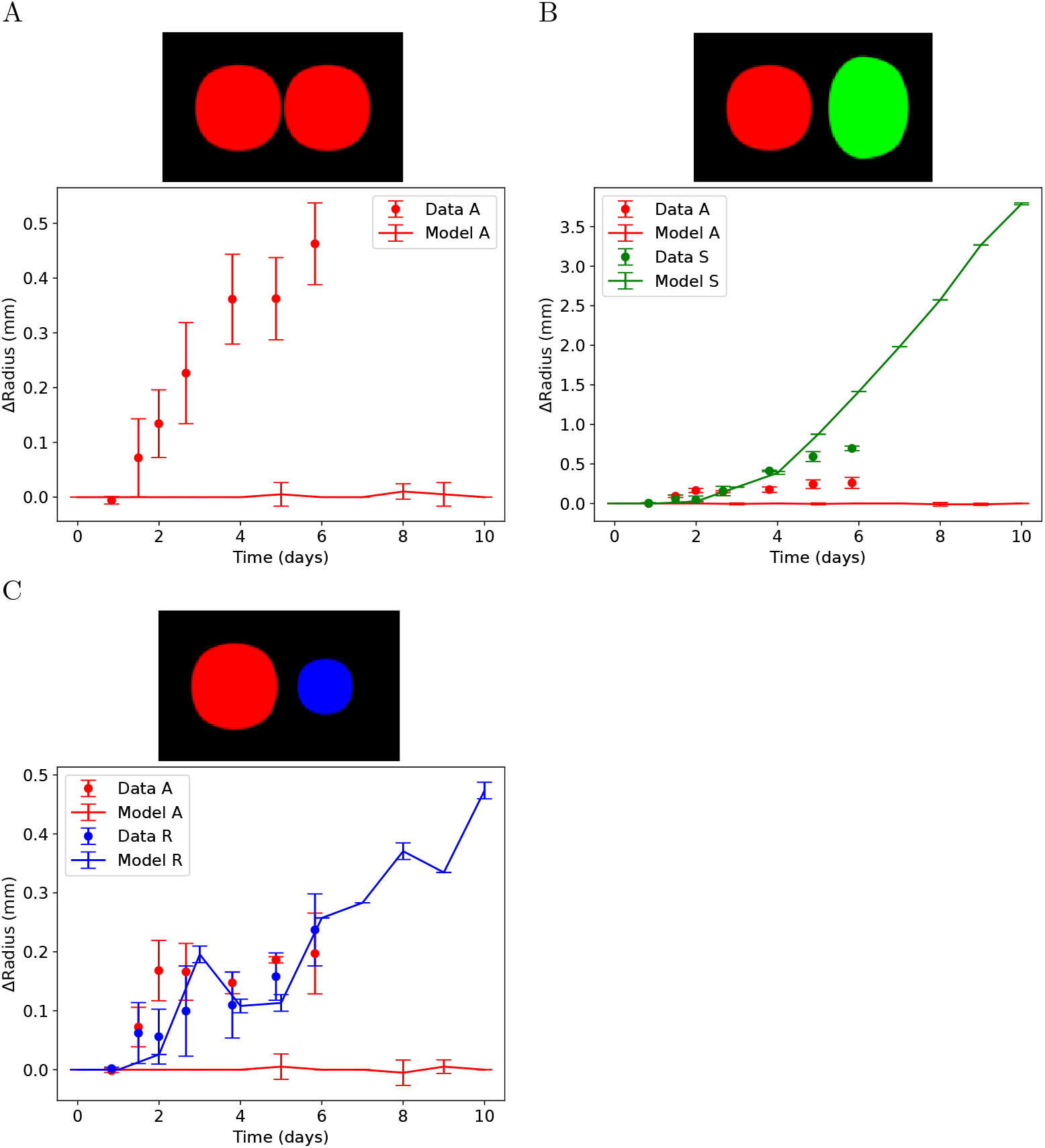
Comparison of the simulations of confronted-colonies experiments with the reported experimental results. ΔRadius is defined as the difference between the colony’s external radius and its internal radius. The top panels present the final state of representative simulations of confronted-colonies experiments, where the antagonistic strain is represented in red, the sensitive strain in green, and the resistant strain in blue. Interested readers may find the corresponding simulations in the repository located at https://github.com/lsanchezg89/facilitationbacterialcommunity.git. The plots depicted below display the average of five simulations, with the error bars indicating the corresponding standard deviation.

As shown in Fig. 4, our model is capable of qualitatively reproducing the observed behavior of sensitive and resistant colonies in the presence of the antagonistic strain. That is, both colonies grow at a slower rate in the direction of the antagonistic colony, albeit for different reasons. While sensitive bacteria die or stop growing as a result of the antagonistic-substance toxicity, resistant bacteria reduce their growth rate as a result of the cost of resisting the antagonistic substance. Nonetheless, in contrast to the findings of Gallardo-Navarro and SantillÁn [44], the antagonistic colony does not show a differential growth rate when confronted with other colonies. This occurs because, according to the model, antagonistic bacteria on the periphery of a given colony not only react to the metabolite (*m*) from bacteria in a nearby colony, but also to that from bacteria within the same colony. We tried several parameter combinations but, because of the above described phenomenon, we couldn’t get a significant difference in the concentration of *m* in the internal and external radii; which is required for differentiated production of antagonistic substance and concomitant growth rate affectation. Because of these drawbacks, we concluded that the mechanisms proposed by Gallardo-Navarro and Santillán [44] are insufficient to explain the observed repertoire of dynamic behaviors. Considering this, we proposed two alternative model amendments: 1) only sensitive and resistant bacteria produce the metabolite sensed by antagonistic bacteria, and 2) all three strains produce metabolite *m*; however, high population density within the colony causes metabolic changes in bacterial cells, resulting in little to no participation in several functions such as replication, metabolite *m* production, or antagonistic substance production.

The first alternative represents the most obvious way to solve the problem of bacteria in the periphery of antagonistic colonies responding to other bacteria within the same colony. However, we know for granted that this model version won’t be able to reproduce the observed growth rate decrease of two facing antagonistic colonies. To account for the fact that antagonistic bacteria do not produce metabolite *m*, the PDE for *m* is modified as follows:

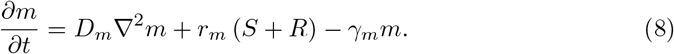

We simulated the confronted colonies experiments with the parameter values in Table 1. The simulation results are summarized in Fig. 5. The simulations corresponding to a couple of facing antagonistic colonies were not carried out for the above explained reasons. Observe that the model is able to qualitatively reproduce the experimental results. Notice in the *A* vs *S* that, in agreement with the experiments, the effect of the antagonistic substance over the sensitive strain is higher than the metabolic cost antagonistic bacteria pay to produce it. Contrarily, the *A* vs *R* simulations predict that the metabolic cost of producing the antagonistic substance *u* is higher than the cost resistant bacteria pay to counteract it. This last outcome disagrees with the experimental observations, where both metabolic costs are comparable.

**Fig 5.**
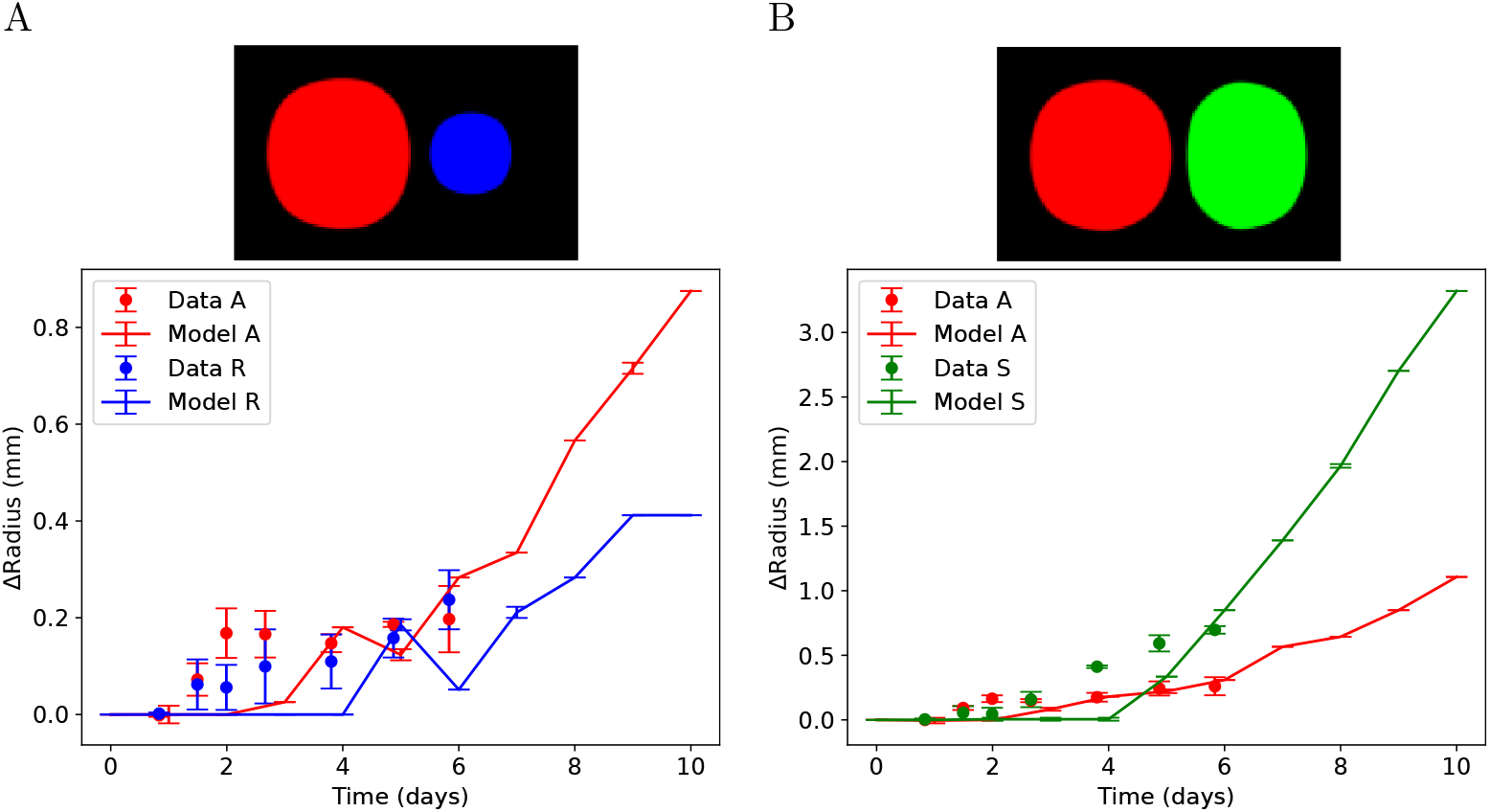
Comparison of the simulations of confronted-colonies experiments with the reported experimental results [44]. ΔRadius is defined as the difference between the colony’s external radius and its internal radius. The top panels present the final state of representative simulations of confronted-colonies experiments, where the antagonistic strain is represented in red, the sensitive strain in green, and the resistant strain in blue. Interested readers may find the corresponding simulations in the repository located at https://github.com/lsanchezg89/facilitationbacterialcommunity.git. The plots depicted below display the average of five simulations, with the error bars indicating the corresponding standard deviation.

To complete the analysis of the first model version, we ran artificial-community simulations with a fixed initial sensitive population and varying initial amounts of antagonistic bacteria, as described in the Methods section. There were two types of simulations: one with no resistant bacteria and another with five times as many resistant bacteria as sensitive bacteria seeded in the simulated area. Gallardo-Navarro and Santillán [44] reported that antagonistic and sensitive bacteria presented competitive exclusion, whereas resistant and sensitive bacteria, and resistant and antagonistic bacteria presented competitive coexistence. Taking this into account, we regarded *α*_*AS*_, *α*_*SA*_, *α*_*RS*_, *α*_*SR*_, *α*_*RA*_ and *α*_*AR*_ (i.e. the Lotka-Volterra competition parameters) as free parameters, and empirically changed their values provided that 0 *< α*_*Rj*_, *α*_*iR*_ *<* 1.0 for *i, j* = *A* or *S* and *α*_*AS*_, *α*_*SA*_ *>* 1.0. The resulting values are tabulated in Tab. 2. After each simulation, we calculated the fraction of the simulated area occupied by sensitive bacteria and plotted it as a function of the initial population of antagonistic bacteria. The results are shown in Fig. 6. In accordance with the experiments, the *S* area-fraction vs. initial-*A*-population curve shifts to the right in the presence of resistant bacteria, implying that resistant bacteria aid sensitive bacteria in surviving the antagonistic strain. Nonetheless, we were unable to obtain results in which one strain (*S* or *A*) drives the other to extinction.

**Fig 6.**
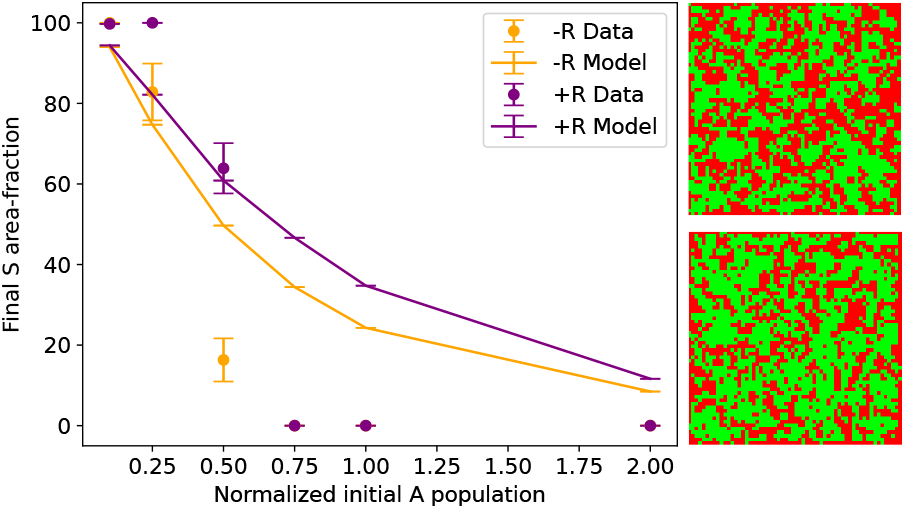
Plots of final area-fraction occupied by the sensitive strain as a function of the initial population of antagonistic bacteria in artificial communities. The reported experimental results [44] are represented with points, whereas the simulation results are plotted with solid lines. Orange points/lines correspond to the experiments in which no resistant bacteria are added to the initial mixture, while purple points/lines correspond to the experiments with added resistant bacteria. Plots represent the average of five simulations and error bars the corresponding standard deviation. The panels on the right-hand side depict the final state of a representative simulated artificial communities. In both of them, the initial condition of antagonistic bacteria (red) is set to 50% of the initial condition of sensitive bacteria (green). The upper panel corresponds to the simulation without the presence of resistant bacteria (not shown), whereas the simulation shown in the lower panel includes the presence of resistant bacteria. The final concentrations were binarized to enhance the contrast. The repository located at https://github.com/lsanchezg89/facilitationbacterialcommunity.git contains the corresponding simulations.

To summarize, assuming that antagonistic bacteria do not produce the metabolite they detect allows the model to qualitatively reproduce many of the experimental results. However, we believe that the outcomes are unsatisfactory, not to mention the model inability to explain antagonistic-bacteria growth-rate decrease in the presence of other antagonistic bacteria. This lead us to analyze a second modification to the original model. Consider that natural selection has established ways of modifying bacteria metabolism to reduce metabolic cost under specific circumstances and when necessary [12, 42]. For example, when bacterial populations reach a high density, bacteria reduce their metabolism and cease participating in processes such as growth, cell division, and metabolite secretion [2, 54–58]. Instead, they begin to act as a group. This behavior benefits the species by making it less vulnerable, and it is also thought to be the key to antibiotic resistance [16, 54]. To incorporate this into the model, we assumed that whenever a local population density is higher than 0.9, the corresponding population remains unchanged as of that moment, stops producing the common metabolite (*m*), and does not interact in any way with bacteria in adjacent compartments. Aside from this consideration, the PDE model remained the same. Parameter estimations required additional fine-tuning in this last model version. The new estimates are tabulated in Table 3.

**Table 3.** Model parameter values for single-colony and confronted-colonies simulations. coefficients than resistant bacteria, in agreement with experimental observations.

| | $A$ | $R$ | $S$ | $m$ | $u$ |
| --- | --- | --- | --- | --- | --- |
| $D$ (mm<sup>2</sup>/min) | $8.5 \times 10^{-9}$ | $5.1 \times 10^{-9}$ | $8.5 \times 10^{-9}$ | 0.51 | 0.085 |
| $r$ (min<sup>-1</sup>) | 0.0078 | 0.0047 | 0.0086 | 0.2 | 0.2 |
| $K$ | 0.004 | $3.5 \times 10^{-15}$ | $0.5 \times 10^{-9}$ | NA | 0.8 |
| $n$ | 3.0 | 3.5 | 3.0 | NA | 3.5 |
| $\gamma$ (min<sup>-1</sup>) | NA | NA | NA | 0.01 | 0.01 |
| $d$ | 0.45 | 0.45 | NA | NA | NA |

Initially, we simulated single colony experiments. The obtained results are summarized in Fig. 7. Again, antagonistic and sensitive bacteria have higher diffusion Next, we simulated the experiments of confronted colonies and presented the results in Fig. 8. Take notice that the model is capable of qualitatively reproducing the experimentally observed behavior in all three situations. That is, all three bacterial strains grow at a slower rate when faced with an antagonistic colony. Nonetheless, the model could not reproduce the experimentally-observed early responses. We only recorded data for 7 simulated days when facing an antagonistic with a sensitive colony because the sensitive colony outgrows the simulated area after that time, making it impossible to measure the external radius.

**Fig 7.**
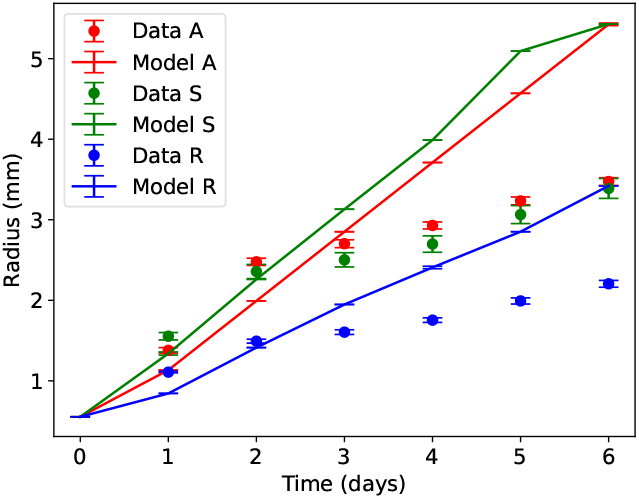
Comparison of the simulations of single colonies growing on agar (solid lines) with and the corresponding reported experimental results (points). Plots represent the average of five simulations and error bars the corresponding standard deviation.

**Fig 8.**
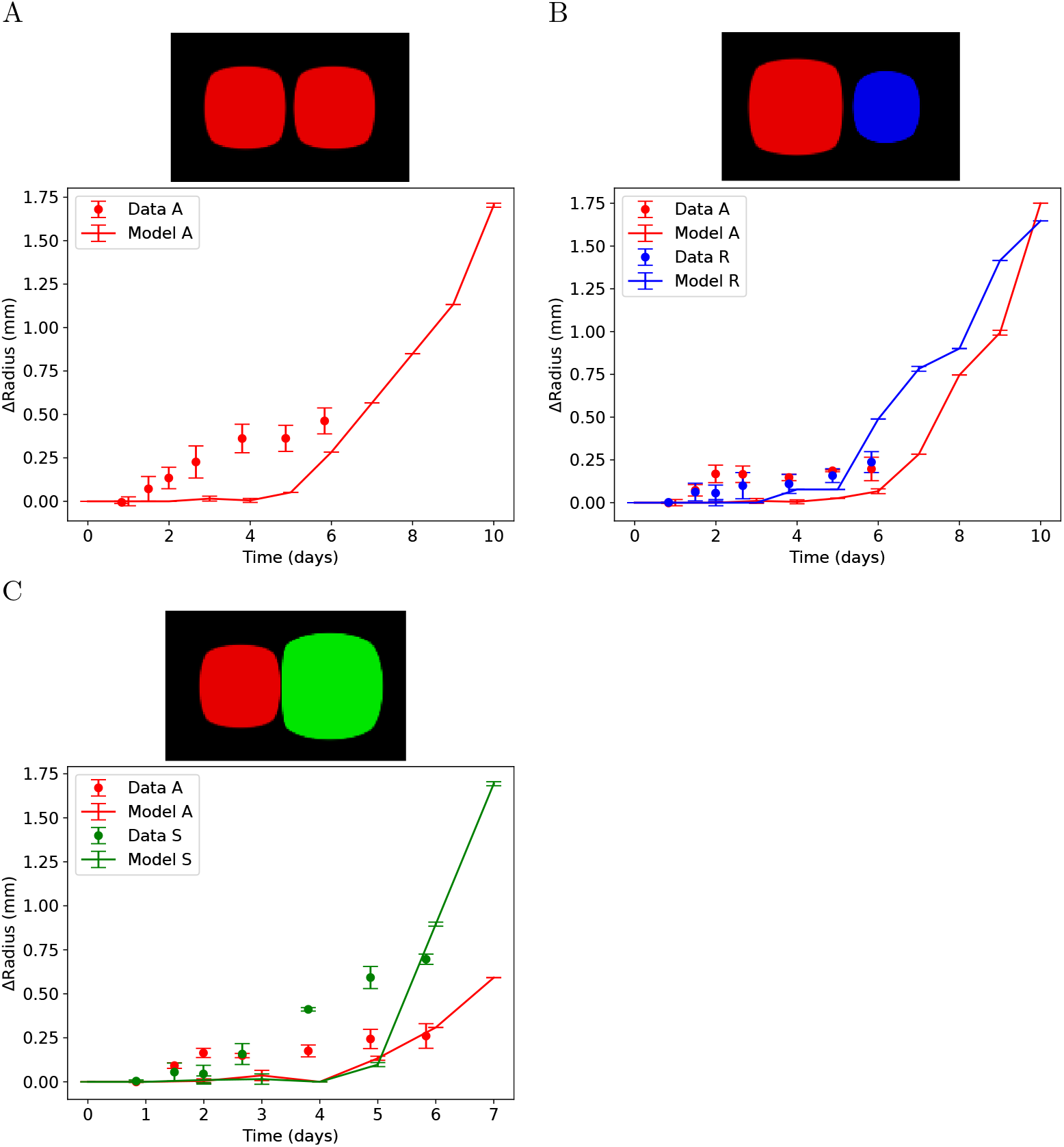
Comparison of the simulations of confronted-colonies experiments with the reported experimental results. ΔRadius is defined as the difference between the colony’s external radius and its internal radius. The top panels present the final state of representative simulations of confronted-colonies experiments, where the antagonistic strain is represented in red, the sensitive strain in green, and the resistant strain in blue. Interested readers may find the corresponding simulations in the repository located at https://github.com/lsanchezg89/facilitationbacterialcommunity.git. The plots depicted below display the average of five simulations, with the error bars indicating the corresponding standard deviation.

In order to further investigate this model version, we simulated the artificial bacterial-community experiments from Gallardo-Navarro and Santillán [44]. After confirming that such simulation times were adequate for the system to reach a steady state, the numerical experiments without resistant bacteria were extended for 10 days, while those with resistant bacteria were extended for 14 days. The close range competition parameters (*α*_*ij*_ with *i, j* = *A, S* or *R*) were re-estimated by fitting the model results to the corresponding experimental data. The new values are listed in table 4. Fig. 9 depicts the simulation results. Take note of how well the model reproduces the reported experimental behavior. In particular, the final *S* area-fraction vs. initial-*A*-population curve shifts to the right in the presence of resistant bacteria, indicating that resistant bacteria help sensitive bacteria survive in the presence of antagonists. Unlike the previous model version, the current one predicts that under certain conditions, the antagonistic and the sensitive strain can drive the other to extinction.

**Table 4.**
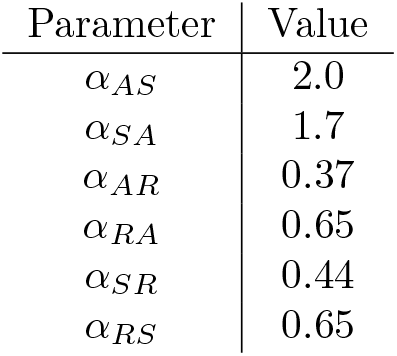
Estimated values for close-range competition parameters, which are employed in the artificial-community simulations.

**Fig 9.**
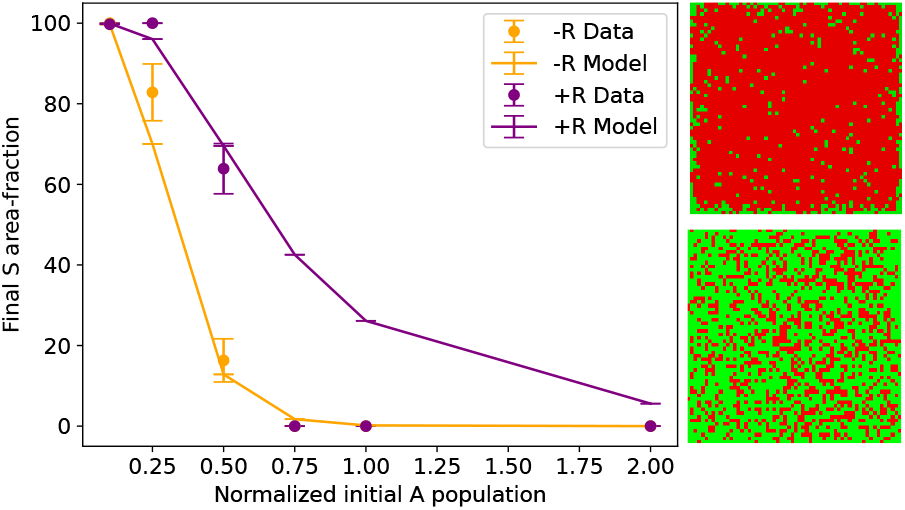
Plots of final area-fraction occupied by the sensitive strain as a function of the initial population of antagonistic bacteria in artificial communities. The reported experimental results are represented with points, whereas the simulation results are plotted with solid lines. Orange points/lines correspond to the experiments in which no resistant bacteria are added to the initial mixture, while purple points/lines correspond to the experiments with added resistant bacteria. Plots represent the average of five simulations and error bars the corresponding standard deviation. The panels on the right-hand side depict the final state of a representative simulated artificial communities. In both of them, the initial condition of antagonistic bacteria (red) is set to 50% of the initial condition of sensitive bacteria (green). The upper panel corresponds to the simulation without the presence of resistant bacteria (not shown), whereas the simulation shown in the lower panel includes the presence of resistant bacteria. The final concentrations were binarized to enhance the contrast. The repository located at https://github.com/lsanchezg89/facilitationbacterialcommunity.git contains the corresponding simulations.

## Discussion and conclusion

We developed a mathematical model to investigate the interaction mechanisms proposed by Gallardo-Navarro and SantillÁn [44] for transfer of resistance against antagonism in a specific set of wild-type bacteria grown in vitro on agar. The community consists of a sensitive, a resistant, and an antagonistic strain, and the proposed mechanisms involve the antagonistic bacteria detecting the proximity of other bacteria through a metabolite (produced by the three of them) that diffuses in the medium. In response, they produce an antagonistic substance that incurs a metabolic cost. The substance kills and inhibits the growth of sensitive bacteria, while resistant bacteria can counteract its effects by incurring a metabolic cost to grow. The presence of resistant bacteria induces the antagonistic ones to produce more antagonistic substance, which decreases their growth rate, and the reduced number of antagonistic bacteria allows the sensitive ones to grow.

According to our results, the mechanisms put forward by Gallardo-Navarro and SantillÁn [44] can account for the toxic impact of the antagonistic substance on sensitive bacteria and the metabolic burden that resistant bacteria experience to overcome the antagonistic substance. However, they fail to account for the metabolic expense that antagonistic bacteria incur to generate the antagonistic substance, since they respond not only to neighboring colonies but also to bacteria within their own colonies. To address this limitation, we proposed the following modifications to the model:

1. Only sensitive and resistant bacteria produce the metabolite sensed by antagonistic bacteria.
2. When high population density is reached within a colony, bacteria undergo metabolic changes and no longer participate in functions such as replication, migration and metabolite or antagonistic substance production.

While the first alternative could explain most of the experimental results in [44], the second alternative provided a more satisfactory explanation. Consequently, we propose that the mechanisms described above are insufficient to clarify how the presence of resistant bacteria can promote the survival of sensitive bacteria. Additional mechanisms are required, such as significant phenotypic changes that occur at high population densities.

Other studies [45, 46] have also reported the role of resistant bacteria in aiding sensitive bacteria to survive in the presence of antagonistic bacteria. Lateral gene exchange is generally invoked as the responsible mechanism. Interestingly, our study supports the possible existence of another mechanism not involving gene exchange that achieves a similar outcome, at least for the specific bacterial community studied by Gallardo-Navarro and Santillán [44], and highlights the significance of spatial distribution in microbial population dynamics. Our findings, along with those from previous studies, suggest that bacteria may have evolved multiple mechanisms to enable the survival of sensitive bacteria in complex communities, despite the various antagonistic interactions present.

The Cuatro Ciénegas valley is an ideal location for studying bacterial communities due to its geography and climate. The Churince lagoon, with its consistent physicochemical properties and the numerous instances of antagonistic interactions among bacterial strains, is a promising site for investigating the effects of biotic interactions. However, conducting studies in this complex ecosystem is challenging, so in vitro studies offer a feasible alternative. Nonetheless, it is essential to note that in vitro conditions differ greatly from in situ, and caution is required when interpreting results. Additional research is necessary to establish the natural occurrence of the mechanism proposed in this study.

The current study is further limited by the use of an explanatory rather than predictive model. It relies on significant simplifying assumptions that make precise simulation of spatiotemporal bacterial dynamics challenging. For example, we assume that the movement of bacteria that causes colonial propagation follows Brownian motion and classical diffusion principles, but empirical evidence suggests that certain types of anomalous diffusion may be more appropriate for these models. Another important simplifying assumption in our model is the use of the logistic model to account for population growth. Generalized logistic models with a competition term based on a power of the population size are known to provide superior fits to experimental data. The intentional use of a simple explanatory model enables the description of the qualitative behavior of the system while requiring a relatively small number of parameters that can be estimated from reported data. However, it is worth noting that this modeling approach limits the quality of fits that can be obtained.

## Acknowledgments

L.S-G. acknowledges CONACyT (México) for her PhD fellowship.

## Notes

### Competing Interest Statement

The authors have declared no competing interest.

